# Assessing jugular venous compliance with optical hemodynamic imaging by modulating intrathoracic pressure

**DOI:** 10.1101/2022.06.28.497963

**Authors:** Robert Amelard, Nyan Flannigan, Courtney A Patterson, Hannah Heigold, Richard L Hughson, Andrew D Robertson

## Abstract

**Significance:** The internal jugular veins are critical cerebral venous drainage pathways that are affected by right heart function. Cardiovascular disease and microgravity can alter central venous pressure (CVP) and venous return, which may contribute to increased intracranial pressure and decreased cardiac output. Assessing jugular venous compliance may provide insight into cerebral drainage and right heart function, but monitoring changes in vessel volume is challenging.

**Aim:** We investigated the feasibility of quantifying jugular venous compliance from jugular venous attenuation (JVA), a non-contact optical measurement of blood volume, alongside CVP from antecubital vein cannulation.

**Approach:** CVP was progressively increased through a guided graded Valsalva maneuver, increasing mouth pressure by 2 mmHg every 2 s until a maximum expiratory pressure of 20 mmHg. JVA was extracted from a 1 cm segment between the clavicle and mid-neck. Contralateral internal jugular vein cross-sectional area (CSA) was measured with ultrasound to validate changes in vessel size. Compliance was calculated using both JVA and CSA between four-beat averages over the duration of the maneuver.

**Results:** JVA and CSA were strongly correlated (median, interquartile range) over the Valsalva maneuver across participants (r=0.986, [0.983, 0.987]). CVP more than doubled on average between baseline and peak strain (10.7 ± 4.4 vs 25.8 ± 5.4 cmH_2_O; *p*<.01). JVA and CSA increased non-linearly with CVP, and both JVA- and CSA-derived compliance decreased progressively from baseline to peak strain (49% and 56% median reduction, respectively), with no significant difference in compliance reduction between the two measures (*Z*=–1.24, *p*=.21). Pressure-volume curves showed a logarithmic relationship in both CSA and JVA.

**Conclusions:** Optical jugular vein assessment may provide new ways to assess jugular distention and cardiac function.

## 1 Introduction

The jugular veins are critical drainage pathways from the head to the heart, with important implications in both cerebral and cardiac health. Cerebral blood drains primarily through the internal jugular veins (IJV) in the majority of healthy individuals in supine position,^1^ although the drainage pathway is dependent on posture and central venous pressure.^2^ Impeded venous return may be reflective of underlying cardiovascular pathology affecting venous return to the right atrium, and may contribute to detrimental long-term effects in cerebral and intracranial health.^3–6^ Thus, assessing jugular venous status may provide insight into cardiac function and cerebral drainage.

Due to the IJV’s high compliance and close proximity to the heart, jugular vein hemodynamics reflect cardiac function and right atrial hemodynamics. In heart failure, the examination of the jugular venous pulse is considered as one of the most important physical markers for determining a patient’s fluid status.^7,8^ However, accurate physical examination of the jugular venous pulse is challenging and highly dependent on clinical experience.^9,10^ Recent studies have investigated the link between jugular vein distention, measured with ultrasound, and congestion.^11,12^ The ratio of IJV size, measured as either cross-sectional area or diameter, between baseline and peak Valsalva strain using ultrasound has been proposed for determining congestion and right atrial status in heart failure.^13,14^ IJV compliance, which describes the vein’s passive volume response to a change in transmural pressure, may provide additional insight into cardiovascular function. Venous compliance traditionally relies on monitoring vessel size as a function of distending pressure. In the limbs, a strain gauge measures tissue volume changes and cannot localize single vessels. Ultrasound can measure cross-sectional vessel area or 3D vessel reconstruction in large vessels, but extreme care needs to be taken by a trained technician to avoid artificially altering the transmural pressure through probe pressure.

Optical technologies have been proposed for monitoring changes in jugular blood volume. Wearable and non-contact optical biosensors have been developed to monitor the jugular venous pulse at various parts of the neck. Reflectance pulse oximeters are able to monitor the jugular venous pulse waveform from the anterior neck.^15^ Additionally, near infrared spectroscopy sensors placed on the lateral neck at the location of the external jugular vein have been shown to predict right atrial pressure and outpatient heart failure risk.^16^ Our previous work has shown that changes in optical attenuation of the jugular vein correlates with changes in central venous pressure (CVP) within the linear range.^17^ Optical technologies provide a promising approach to non-intrusively monitor changes in jugular vein volume, but they have not been investigated for quantifying jugular compliance across a wide range of pressures.

In this paper, we sought to quantify the jugular venous compliance curve by relating changes in vessel size, using non-contact optical hemodynamic imaging, to CVP from antecubital cannulation across a wide range of pressures. We hypothesized that optical attenuation from the jugular vein would reflect the underlying vessel size across the non-linear range of pressure values, evaluated using a graded Valsalva maneuver to increase intrathoracic pressure, and thus CVP.

## 2 Methods

### 2.1 Participants

This study was conducted as part of a larger study for which data collection protocol and recruitment were previously presented.^17^ Briefly, 9 young healthy adults (5 female, age 25.4 ± 5.6 years, height 166.2 ± 11.4 cm, weight 63.8 ± 12.8 kg; BMI 22.9 ± 2.2 kg m^-2^) with no history of vascular, inflammatory, or thrombotic disease were included in this study. Participants were asked to fast for at least 2 h and consume 1 L of water 2 h prior to testing, as well as to abstain from caffeine, alcohol consumption, and heavy exercise for 24 h prior to testing. The study was approved by a University of Waterloo Research Ethics Committee (ORE #40394) and all protocols conformed to the principles of the Declaration of Helsinki. Participants provided written informed consent prior to testing.

### 2.2 Data Collection and Experimental Protocol

Participants were instrumented with an electrocardiogram (ECG; iE33 xMatrix, Philips Health-care, Andover, USA) and continuous arterial blood pressure finger plethysmogram with estimated stroke volume (SV) via Modelflow (NOVA, Finapres Medical Systems, Enschede, NL). Brachial artery blood pressures, which are reported here, were reconstructed using those from the finger artery and calibrated to arm cuff pressures (NOVA). Cerebral blood velocity was monitored by insonating the left middle cerebral artery by transcranial Doppler ultrasound using a 2 MHz transducer (Neurovision 500M, Multigon Industries, Elmsford, USA). Continuous CVP was measured using a catheter placed in a right antecubital vein.^18^ The vein was punctured using a single-use 20G needle and attached to a pressure monitoring transducer through a saline-filled polyurethane catheter. Participants were supine and rightward-tilted to ensure a continuous column of blood between the right atrium and the antecubital vein. The pressure transducer was calibrated at 0, 5, 10, and 15 cmH_2_O relative to the right midclavicular line using a laser level. Simultaneous vascular ultrasound of the left IJV, in cross-section, was conducted using B-mode ultrasound with a 9-3 MHz linear transducer (iE33 xMatrix).

The optical imaging system was previously presented in detail.^17^ Briefly, tissue was illuminated with a 940 nm near-infrared LED (LZ1-10R702, LED Engin, San Jose, USA) passed through a mounted diffused spot beam lens optic, custom uniform beam optic, and near-infrared polarizer. The optical imaging system was positioned such that it provided orthogonal illumination^19^ of the right lateral area of the neck. Tissue reflectance was cross-polarized to minimize surface-level specular reflection, and imaged by a near-infrared sensitive camera (GS3-U3-41C6NIR-C, Teledyne FLIR, Wilsonville, USA). A near-infrared bandpass filter was mounted in front of the camera lens to minimize ambient visible spectrum light. The components were integrated into a 3D printed enclosure to fix the orientation of the LED assembly and camera optics. A time-synchronized arterial signal was captured through a finger photoplethysmography sensor to identify the locations of the pulsatile venous signal (see Section 2.3). The imaging system acquisitions were time-synchronized with signal acquisition through a hardware trigger to the acquisition board.

CVP was altered by changing the intrathoracic pressure using a guided graded Valsalva protocol. Following a 15 s baseline period of self-paced breathing, participants began forced obstructed expiration through a tube in their mouth that was connected to an external digital pressure manometer. Participants were audibly instructed to increase their expiratory pressure by 2 mmHg every 2 s until a maximum of 20 mmHg (27.2 cmH_2_O). Participants matched the stated target pressure through visual feedback from the manometer.

Although participants were instructed on generating changes in intrathoracic pressure with a practice maneuver, it is nonetheless possible that the pressures could be achieved through buccal maneuvers, especially at the lower initial pressures. Thus, a small hole was punctured in the mouth-piece so that participants exerted continuous expiratory pressure from the thorax.^20^ The respiratory and cardiovascular signals were visually evaluated to confirm that proper Valsalva maneuvers were achieved.

### 2.3 Data Processing

#### 2.3.1 Jugular Venous Attenuation

In the case of a distending vessel during a Valsalva maneuver, changes in optical attenuation are proportional to the increased mean photon path length through the vessel:^17^

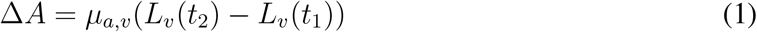

where *μ_a,v_* and *L_v_* are the absorption coefficient and mean path length through the vessel. Thus, changes in vessel size could be monitored by quantifying optical attenuation at the location of the jugular vein.

The details of computing a jugular venous pulsatility map for extracting a jugular pulse wave-form were previously presented.^17^ Briefly, distinct signal locations were computed from the diffuse reflectance frames using a 4×4 pixel mean filter, and each location’s temporal signal (“virtual sensor”) was denoised using a hemodynamic-inspired Kalman filter.^21^ These denoised diffuse reflectance frames were converted to optical attenuation frames, where each virtual sensor signal encoded the location’s dynamic attenuation modulated by hemodynamic processes. Motion induced by respiratory effort was corrected through a dual correlation filter^22^ using a linear kernel with histogram of gradient features by selecting a small salient region of interest (e.g., skin lesion, surface vessel) in the grayscale reflectance frames. Noting that arterial and jugular pulse waveforms are out of phase,^23^ the locations of the venous pulse were identified by computing a venous pulsatility map through a temporal negative correlation filter relative to the time-synchronized arterial waveform, and negatively correlated values were visually coded and displayed over the original reflectance frames. The venous pulsatily map was used to extract a jugular venous attenuation (JVA) wave-form as the average response across an approximately 1 cm long region within the center of the vessel between the clavicle and mid-neck.

#### 2.3.2 Beat-by-beat extraction

ECG and blood pressure signals were recorded at 1000 Hz (PowerLab with LabChart v7.3.7; ADInstruments, Colorado Springs, CO), and optical imaging was captured in uncompressed video format at 60 fps with 16 ms exposure time. These signals were time-synchronized at 60 Hz for processing. The signals were analyzed in LabChart and MATLAB 2020b (MathWorks, Natick, MA). Heart rate (HR), mean arterial pressure (MAP), SV, mean CVP, mean JVA, and mean cerebral blood velocity were extracted as beat-by-beat variables between consecutive ECG R waves. Cardiac output was calculated as the product of HR and SV. Ultrasound imaging was acquired at approximately 23 fps. The first frame following each ECG R wave was extracted for quantification of IJV cross-sectional area (CSA). CSA was measured semi-automatically using the ellipse tool in ImageJ and adjusted manually as necessary to fit the lumen-wall border.

#### 2.3.3 Venous Compliance

Four-beat averages were used in compliance computations to account for small inter-beat variations that may cause negative compliance values at low pressures and close to zero pressure changes. Baseline values were computed as the 4-beat average starting 10 s prior to Valsalva. In the cases where the end of Valsalva was reached before a fourth beat, the last complete four-beat sequence was used. Four-beat average differences of the variables of interest were computed using the central difference method:

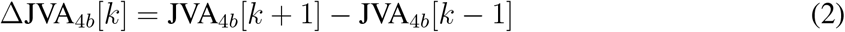

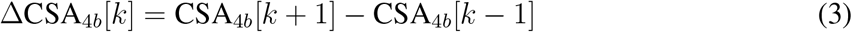

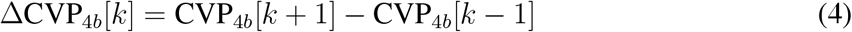

Venous compliance values were calculated across a range of CVP attained through the guided graded Valsalva protocol (see Section 2.2). Compliance is generally defined as a change in volume over a change in pressure. The change in pressure was attained through modulating CVP. Two variables were used as proxies for changes in volume: change in JVA (optical) and change in CSA (ultrasound).

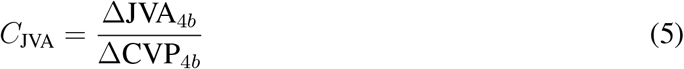

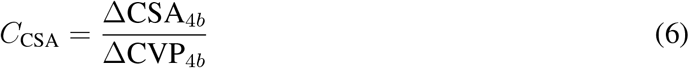

CSA-derived compliance was used to validate observations of JVA-derived compliance.

### 2.4 Statistical Analysis

Statistics were computed in R 4.1.2.24 The per-participant relationship of the four-beat average CSA and JVA were compared by fitting a linear regression model, and linear correlation coefficients are reported. Paired sample t-tests were used to compare baseline and peak Valsalva responses. Normality of the data was confirmed by the Shapiro-Wilks test. Wilcoxon signed rank test was used to compare paired differences in baseline-to-peak compliance reduction between *C*_CS_A and *C*_JVA_. We reported statistically significant results when *p* < .05. Data are reported as mean ± SD.

## 3 Results

Figure 1 shows the group responses of CVP, CSA, and JVA to the graded Valsalva protocol. Mean baseline CVP was 10.7 ± 4.4 cmH_2_O in supine posture (Table 1). During the Valsalva strain, CSA and JVA increased with CVP at low pressures. At higher pressures, however, CVP increases faster than both CSA and JVA, indicating reduced jugular compliance. Four-beat average JVA and CSA were strongly correlated (median, interquartile range) across all participants (r=0.986, [0.983, 0.987]) (see Figure 2).

**Fig 1:**
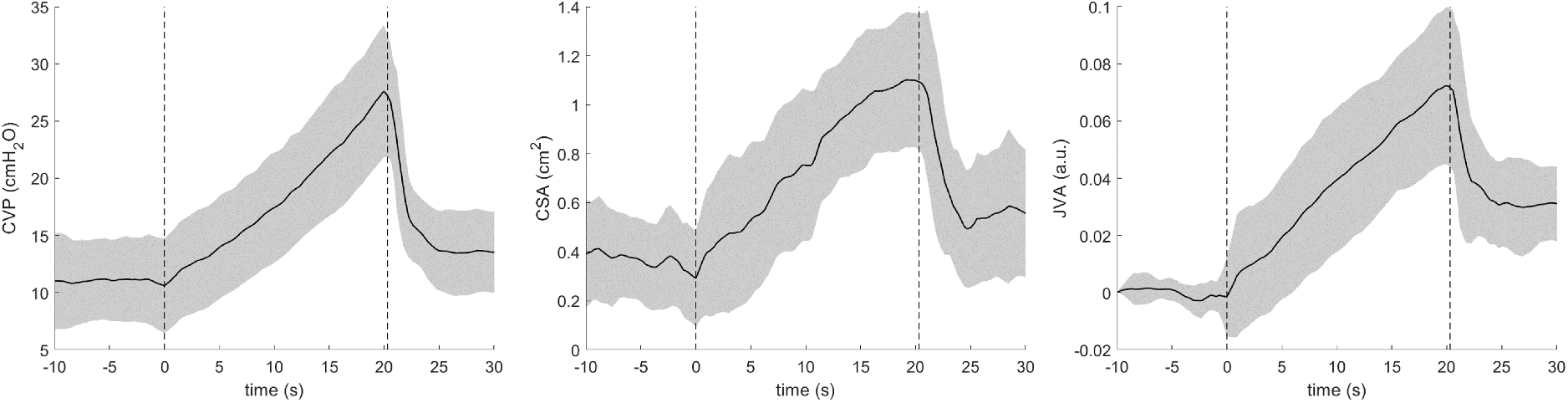
Group responses to graded Valsalva maneuver. Stepwise (per 2 s) increases of 2 mmHg in mouth pressure translated to progressive increases in central venous pressure (CVP), internal jugular vein cross-sectional area (CSA), and optical jugular venous attenuation (JVA) relative to baseline. Dashed lines represent the start/stop of Valsalva strain. Shaded areas represent ±SD.

**Fig 2:**
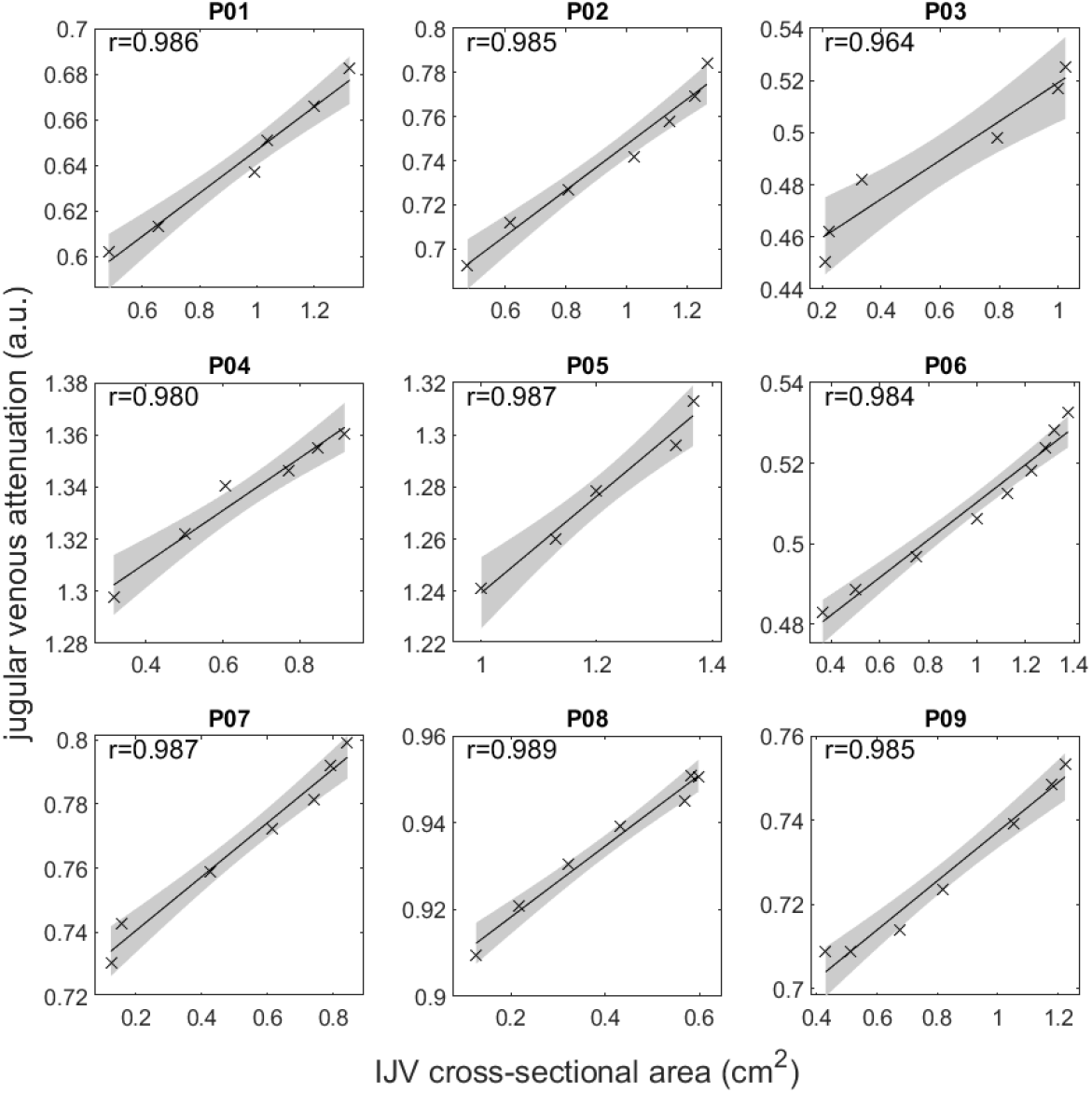
Individual correlation plots between internal jugular vein (IJV) cross-sectional area and jugular venous attenuation four-beat averages across the graded Valsalva maneuver.

**Table 1:** Cardiovascular variables assessed at rest and peak Valsalva strain while supine.

|  | Baseline | Peak Valsalva | *p* value |
| --- | --- | --- | --- |
| Heart rate, beats/min | 67.9 ± 12.8 | 79.7 ± 17.9 | .04 |
| Stroke volume, ml | 81 ± 14 | 55 ± 13 | <.01 |
| Cardiac output, l/min | 5.4 ± 1.1 | 4.2 ± 0.7 | .01 |
| Systolic blood pressure, mmHg | 119.4 ± 9.0 | 125.7 ± 17.2 | .11 |
| Diastolic blood pressure, mmHg | 71.1 ± 5.7 | 85.6 ± 10.0 | <.01 |
| Mean arterial pressure, mmHg | 92.7 ± 4.6 | 104 ± 12.4 | .02 |
| Central venous pressure, cmH<sub>2</sub>O | 10.7 ± 4.4 | 25.8 ± 5.4 | <.01 |
| Internal jugular vein cross-sectional area, cm<sup>2</sup> | 0.4 ± 0.2 | 1.1 ± 0.3 | <.01 |
| Jugular venous attenuation, a.u. | 0.78 ± 0.29 | 0.85 ± 0.30 | <.01 |
| Mean cerebral blood velocity, cm/s | 75.3 ± 15.7 | 76.3 ± 17.4 | .70 |

Systemic and local cardiovascular variables were significantly altered at peak Valsalva strain compared to resting supine baseline (Table 1). Central venous pressure more than doubled on average between baseline and peak strain (10.7 ± 4.4 vs 25.8 ± 5.4 cmH_2_O; *p* < .01). This combined with a 31% mean reduction in stroke volume indicated impeded venous return, with commensurate increases in both CSA (0.4 ± 0.2 vs 1.1 ± 0.3 cm^2^; *p* < .01) and JVA (0.78 ± 0.29 vs 0.85 ± 0.30 a.u.; *p* < .01). Stroke volume was reduced (81 ± 14 vs 55 ± 13 ml; *p* < .01), and despite an increase in heart rate (67.9 ± 12.8 vs 79.7 ± 17.9 beats min^-1^; *p* = .04), cardiac output fell (5.4 ± 1.1 vs 4.2 ± 0.7 1 min^-1^; *p* < .01). Mean arterial pressure increased significantly, and although mean cerebral blood velocity was unchanged across the sample, cerebral blood velocity increased in all but two of the participants (Figure 3; see Discussion).

**Fig 3:**
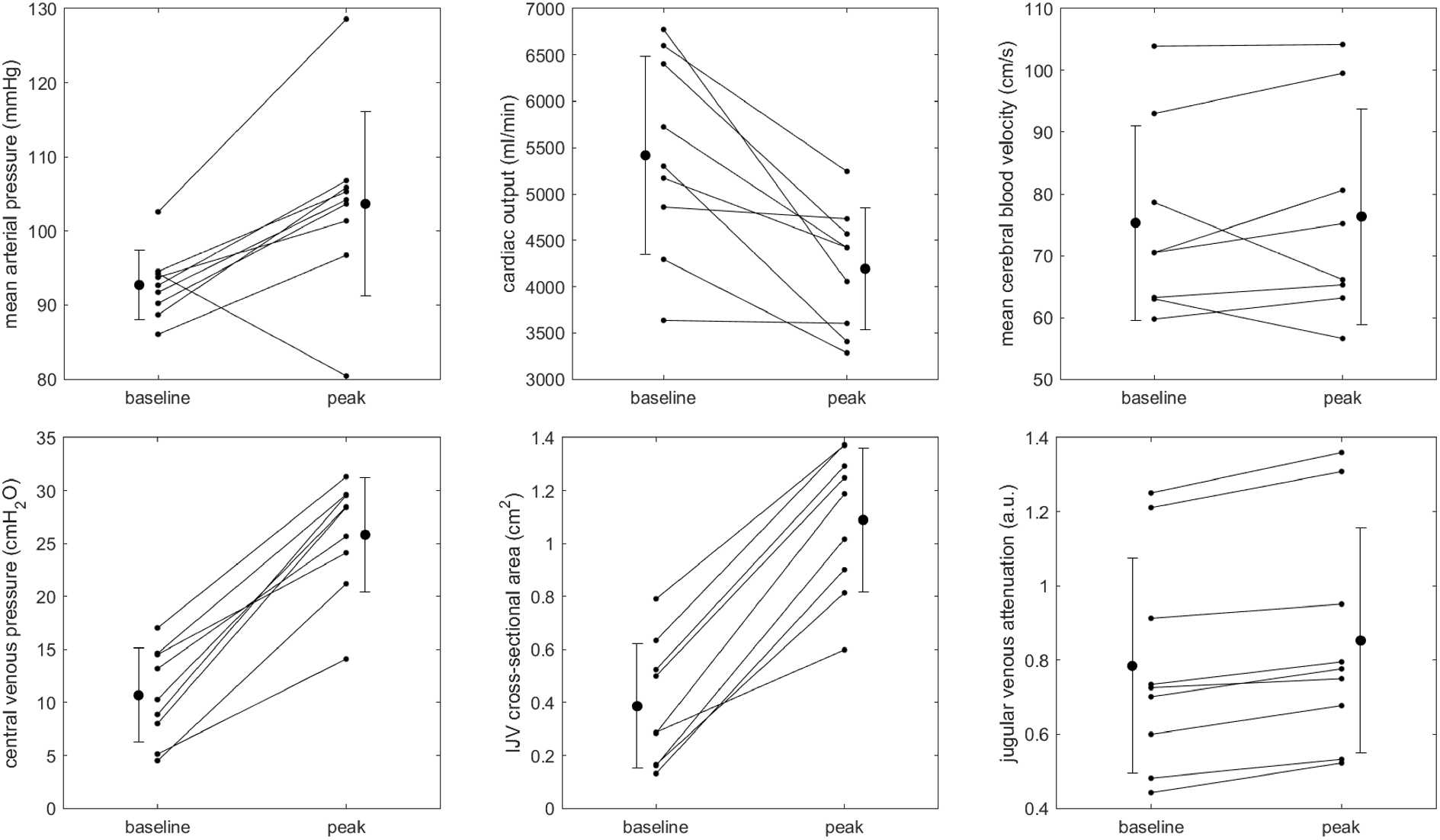
Individual data showing changes in cardiovascular variables from supine baseline to peak Valsalva strain at 20 mmHg. All variables except mean cerebral blood velocity significantly increased from baseline to peak (see Table 1 for statistical analyses). Error bars represent ±SD.

Individual pressure-volume curves show a logarithmic relationship between pressure and volume in both CSA and JVA data (Figure 4), indicating a non-linear reduction in compliance as pressure increased. Baseline CVP ranged from 5.7 to 17.9 cmH_2_O. Figure 5 shows individual compliance curves derived from each participant’s pressure-volume curves. As CVP increased, both JVA-derived (*C*_JVA_) and CSA-derived (*C*_CSA_) compliance decreased, and the rate of decrease was reduced at higher pressures. The data demonstrate large changes in volume for small changes in pressure at the initial low pressures (*C*_JVA_: (8.1 ± 7.1) × 10^-3^ a.u. cmH_2_O^-1^; *C*_CSA_: (8.4 ± 7.3) × 10^-2^ cm^2^cmH_2_O^-1^), and a slowed reduction in compliance at high pressures as the vein converged toward maximal distention (*C*_JVA_: (3.1 ± 0.1) × 10^-3^ a.u.cmH_2_O^-1^; *C*_CSA_: (2.4 ± 0.8) × 10^-2^ cm^2^cmH_2_O^-1^), resulting in median compliance reduction of 49% and 56% respectively (Figure 4b). The percent reduction in compliance between both methods were not significantly different (*Z*=–1.24, *p* = 0.21).

**Fig 4:**
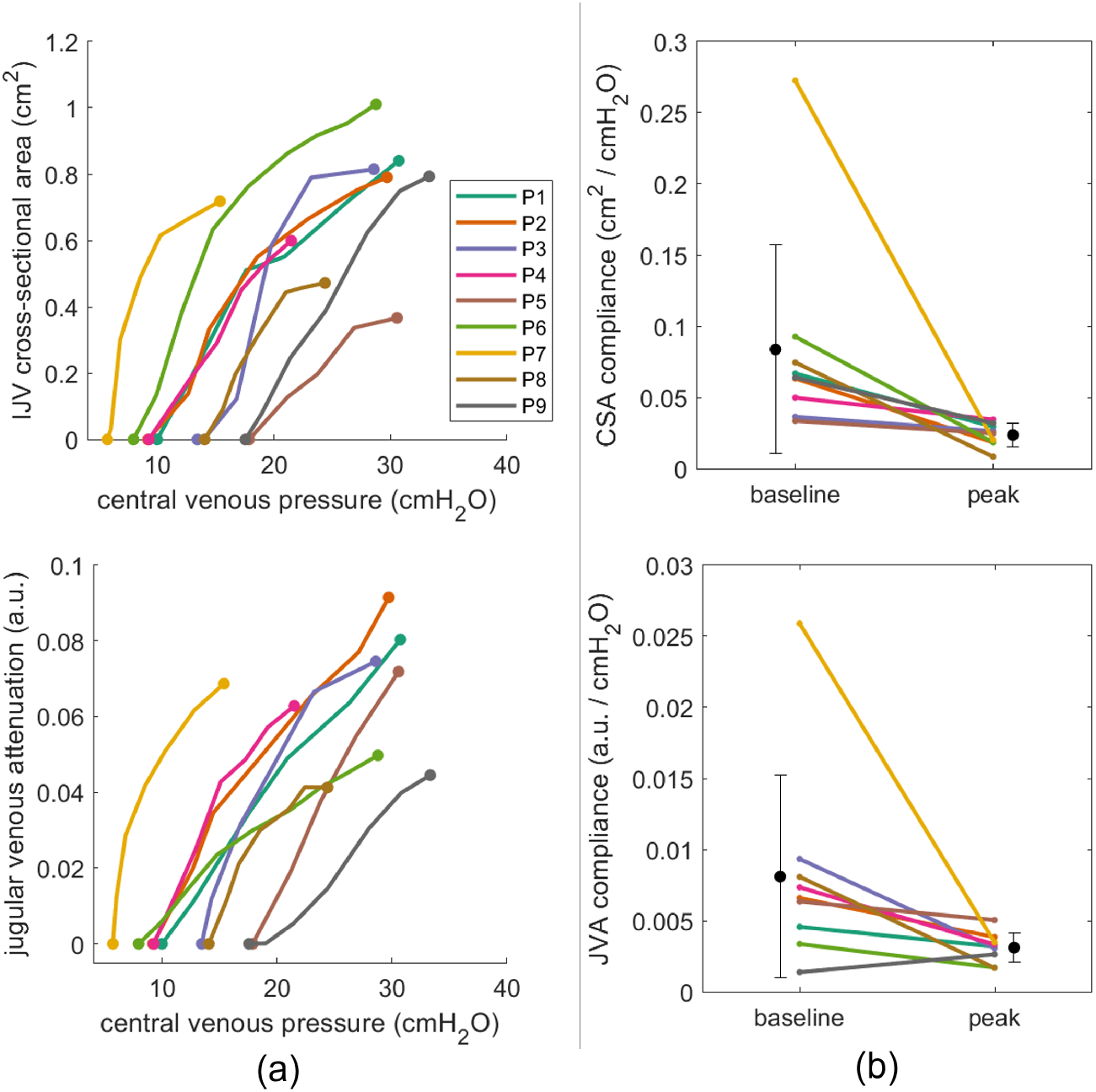
Individual pressure-volume curves (a) and compliance changes (b) during graded Valsalva from 0 to 20 mmHg over 20 s, using internal jugular vein (IJV) cross-sectional area (top row) and normalized jugular venous attenuation (bottom row) as proxies for changes in vessel volume.

**Fig 5:**
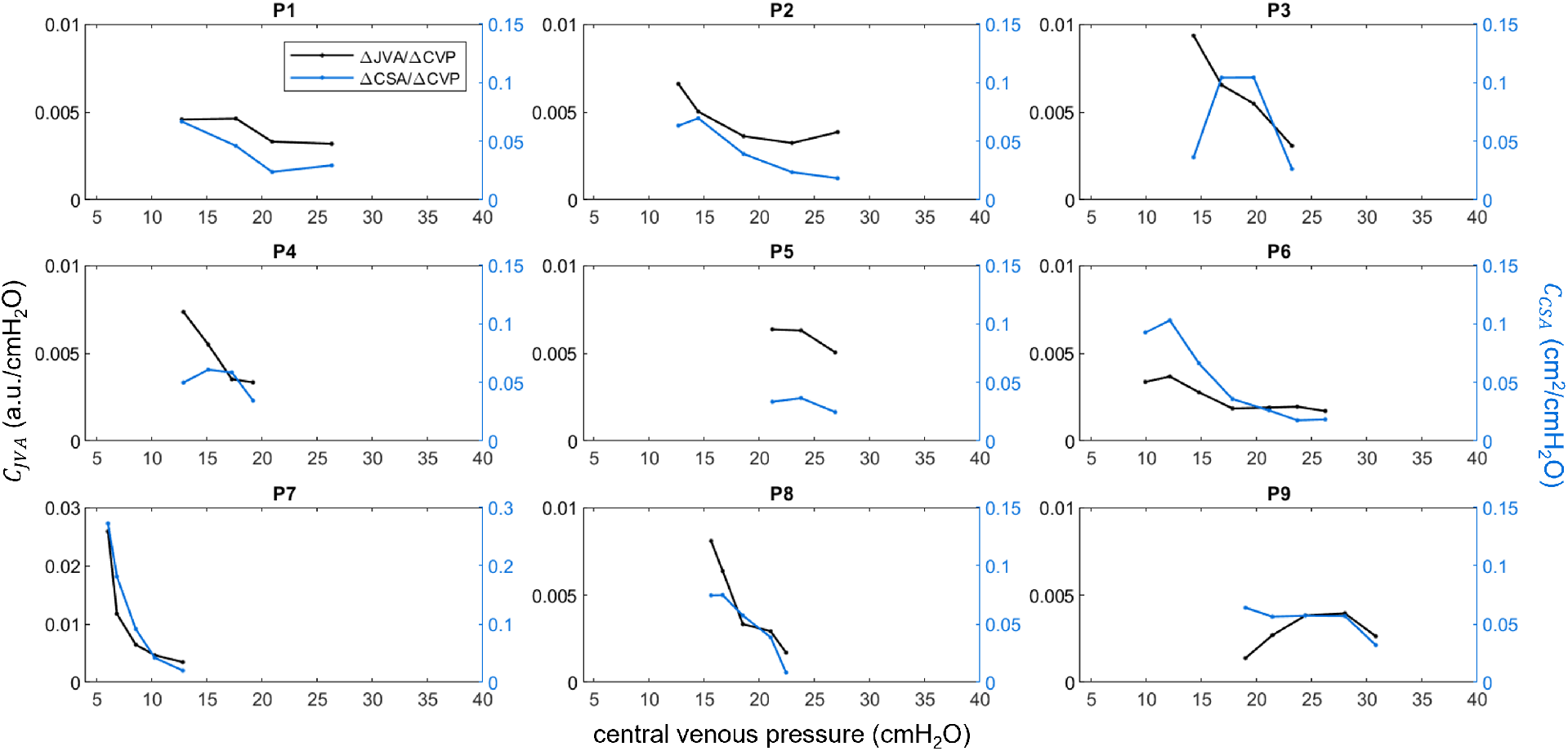
Individual compliance curves calculated from each participant’s pressure-volume curves using both jugular venous attenuation (JVA; black) and internal jugular vein cross-sectional area (CSA; blue) as proxies for vessel volume. Compliance decreased as central venous pressure (CVP) increased during graded Valsalva maneuver. Each point represents a four-beat average during the graded Valsalva maneuver.

In the case with the lowest resting CVP (P7, bottom left panel in Figure 5), a rapid initial drop in compliance was observed, indicating a very compliant vessel at baseline. Note that the y-axis for this participant was increased to accommodate the initial high compliance.

## 4 Discussion

In this study, we applied a progressive increase in intrathoracic pressure through a Valsalva maneuver to gain insight into jugular vein distension and compliance. Previous studies have proposed ultrasound metrics of vessel size changes from baseline to peak of a Valsalva maneuver to determine congestion and right atrial status in heart failure.^13,14^ A smaller increase in diameter is a direct consequence of elevated right atrial pressure with an already partially distended jugular vein during rest. Our data, obtained in healthy young adults, showed that JVA by optical imaging can be an accessible non-contact alternative to ultrasound to assess jugular vein compliance and right heart status. The compliance curves derived from JVA may provide additional insight into venous status using a controlled graded intrathoracic pressure change.

We observed a strong linear correlation between CSA and JVA, indicating that JVA tracked changes in volume well when considering CSA as a proxy for vessel volume, and expanding on our previous findings that JVA tracks changes in CVP within the linear range,^17^ as well as CSA across head-down tilt and lower body negative pressure provocations.^25^ Calculating compliance using CSA (cross-sectional compliance^26^) assumes that changes in vessel volumes and vessel diameters are linearly correlated. By calculating compliance at a particular location along the jugular track, we demonstrated that optical imaging provides non-intrusive jugular venous compliance assessment for a unit vessel length. As the IJV has variable shape along the length of the vessel,^27^ expanding this analysis across the length of the neck may further advance insights into longitudinal distention and filling patterns.

Since JVA is a measure of optical attenuation, which is expressed in arbitrary units, *C*_JVA_ is expressed in a.u. cmH_2_O^-1^. Although we have observed a strong linear correlation between JVA and CSA across pressures and general agreement between *C*_CSA_ and *C*_JVA_, it becomes challenging to compare compliance values between individuals due to differences in endogenous tissue chromophores and scattering agents.^28^ However, the ratio between maximum jugular vein diameter during Valsalva to diameter at rest, and not maximum jugular vein diameter alone, was found to be a significant risk factor for heart failure hospitalization or mortality.^13^ Thus, in the context of right atrial screening, tracking changes within an individual is more important than comparing absolute compliance values between individuals.

Compliance changes when CVP was above 20 cmH_2_O (14.7 mmHg) were marginal across participants that attained this pressure. In our protocol, participants performed the Valsalva while supine, thus there were no hydrostatic effects between the jugular vein and the heart, so CVP reflected right atrial pressures. This observed effect is consistent with clinical guidelines on assessing congestion thresholds, which define elevated congestion at 15 mmHg (20.4 cmH_2_O) in transthoracic echocardiography and pulmonary hypertension.^29,30^ Patients with small jugular vein diameter ratios in a Valsalva maneuver are more likely to have signs of congestion.^13^ Further investigations are needed to determine whether this asymptotic compliance pattern may be suitable for assessing congestion in clinical populations.

Increased MAP, CVP, and cerebral blood velocity in 7 of 9 participants suggests that intracranial pressure increased from baseline to peak Valsalva, consistent with previous findings.^31^ Venous outflow is an important contributor to intracranial pressure regulation.^32^ Paired with MAP, changes in JVA may indicate impaired cerebral drainage secondary to elevated CVP. Although the Valsalva maneuver provides a convenient way to assess acute pressure-induced sympathoexcitatory response while increasing right atrial pressure, further investigations are needed to assess the feasibility of longitudinal monitoring of cerebral drainage for applications such as monitoring intracranial hypertension^33^ and remodeling of the eye in spaceflight.^5,6^

Although individual compliance curves showed a general decrease in compliance as CVP increased, a transient rise in *C*_CSA_ was observed at low pressures (approximately 15 cmH_2_O) in some participants. This was not observed in the corresponding *C*_JVA_ curves, and after these initial pressures, *C*_CSA_ and *C*_JVA_ curves matched each other. Given these empirical observations, these observed transient increases may have been a result of small ultrasound probe hold-down pressure restricting vessel distention at low CVP. At 15 cmH_2_O (11 mmHg), a probe pressure of 0.15 Ncm^-2^ (0.21 psi) is sufficient for collapsing the jugular vein. This provides motivation for non-contact approaches for quantifying vessel distention when low pressures are of interest.

We observed a decrease in mean arterial pressure from baseline to peak Valsalva strain in one female participant (P7 in figures above). Consequently, mean cerebral blood velocity decreased. This participant had the lowest resting CVP and fastest initial change in compliance, experienced transient symptoms (hearing disturbance and lightheadedness) during a separate expiratory effort of 40 mmHg and extremity numbness during −40 mmHg lower body negative pressure (data not shown), and did not consume food or water for at least 8 h before arriving at the lab, despite the instructions to consume 1 L of water prior to the study. We therefore hypothesize that the participant was hypovolemic, causing low cardiac filling pressures and plasma volume, which has been associated with marked reductions in arterial blood pressure during Valsalva strain.^34^

Non-contact optical monitoring of the jugular vein may provide new ways for assessing cardiovascular health in individuals affected by changes in fluid status, such as in heart failure, sepsis, and spaceflight. Clinically relevant jugular physiology, such as distention^7,35^ and stasis,^36^ is often difficult to discern visually.^9,10^ Volume-related biomarkers are less susceptible to cardiovascular compensatory mechanisms, and thus may provide earlier detection of fluid-related pathophysiology.^37^ Optical assessment of jugular venous compliance may be helpful in identifying jugular venous abnormalities secondary to changes in cardiac and fluid status.

There are limitations to this study. The sample size was small, thus group statistics may not be appropriate for other populations. The participants were healthy young adults, thus further studies are needed to investigate jugular venous dynamics in other populations (e.g., clinical, obese, older adults, and pediatrics).

## 5 Conclusion

In this paper, we investigated the feasibility of assessing dynamic jugular venous compliance using non-contact optical imaging measures of vessel volume changes in a healthy young adult sample. Compliance was calculated over a range of pressures attained by modulating the intrathoracic pressure through a graded Valsalva maneuver. Jugular venous attenuation (JVA; optical) was compared to internal jugular vein cross-sectional area (CSA; ultrasound) as vessel volume proxies for calculating pressure-volume and compliance data. Changes in JVA and CSA were strongly correlated during graded Valsalva across participants. JVA significantly increased between baseline and peak strain, with a commensurate significant increase in CSA and central venous pressure. Pressure-volume curves, using JVA and CSA as proxies for vessel volume, showed logarithmic volume responses as pressure increased. Both JVA- and CSA-derived compliance decreased as pressure increased, with a slowed reduction at high pressures as the vein converged toward maximal distention. These results showed that JVA was able to quantify changes in IJV volume without the potential for compression at low venous pressures. Taken together, these results show promise for using non-contact optical imaging to assess dynamic changes in vessel volume for evaluating congestion in heart failure and venous biomarkers secondary to altered cardiac and fluid status.

## Acknowledgments

This work was supported by the Natural Sciences and Engineering Research Council of Canada (PDF-503038-2017) and Canada Space Agency (18FAWATA13).

## Code, Data, and Materials Availability

In accordance with the ethical requirements of human data handling, participant data cannot be made publicly available. Anonymized processed data can be made available upon reasonable request.

